# Microinjection in *C. elegans* by direct penetration of elastomeric membranes

**DOI:** 10.1101/2022.05.11.491555

**Authors:** Shawn R. Lockery, Stelian Pop, Ben Jussila

## Abstract

The nematode worm *C. elegans* is widely used in basic and translational research. The creation of transgenic strains by injecting DNA constructs into the worm’s gonad is an essential step in many *C. elegans* research projects. This paper describes the fabrication and use of a minimalist microfluidic chip for performing microinjections. The worm is immobilized in a tight-fitting microchannel one sidewall of which is a thin elastomeric membrane which the injection pipette penetrates to reach the worm. The pipette is neither broken nor clogged by passing through the membrane, and the membrane reseals when the pipette is withdrawn. Rates of survival and transgenesis are similar to those in the conventional method. Novice users found injections using the device easier to learn than the conventional method. The principle of direct penetration of elastomeric membranes is readily adaptable to microinjections in a wide range of organisms including cells, embryos, and other small animal models. It could therefore lead to a new generation of microinjection systems for basic, translational, and industrial applications.

## Introduction

The nematode *Caenorhabditis elegans* is a preeminent research organism in biology and medicine. Its small size, ease of culture, rapid development, and large mutant libraries make this organism a powerful model for assigning functions to genes relevant to human disease^1^. These strengths are paired with unsurpassed genetic tractability, including a prodigious molecular biological toolkit and the most comprehensively annotated genome to date.

Transgenesis – the transfer of DNA into the germline of an organism – is an essential step in many *C. elegans* research projects. This critical technology has changed very little since its introduction more than 30 years ago^2^. Conventional *C. elegans* transgenesis involves four main steps: (i) Manually mounting up to 10 worms on a dry agarose pad formed on cover slip; (ii) Transferring the coverslip to a compound microscope fitted with a micromanipulator and pipette holder to position the injection pipette; (iii) Inserting the pipette tip into the gonad of each worm and injecting a solution of DNA; (iv) Manually recovering each injected animal from the coverslip to a standard culture plate. In this procedure, the physical resistance of the worm to penetration by the pipette critically depends on the fact that the agarose substrate is dry. The drawback of this method of immobilization is that it rapidly desiccates the animal. To obtain reasonable survival rates, this procedure therefore must be performed quickly, in less than 10 min, or about 1-2 minutes per worm. The need for both speed and dexterity makes the technique difficult to master and tiring to perform; even experienced investigators rarely inject more than four to six strains per day. There is therefore a need for transgenesis methods that are easier to learn and less tiring.

In response to this need, at least six alternative nematode injection systems have been validated at the level of proof of concept^3–8^. Each of these systems seeks to eliminate the main obstacle to facile DNA injections: fixing the animal in place firmly enough to permit penetration by the injection pipette but without desiccation. In these systems, the worm is fixed in a wet environment one of two ways: by encapsulation in a temperature sensitive hydrogel^4^, or by entrapment in a customized, fluid-filled microfluidic device^3,5–8^. A common feature of all six microfluidic systems is the establishment of an unobstructed pathway by which the injection pipette reaches the worm. These systems fall into two main categories: (i) *Open systems*, in which the compartment holding the worm has no ceiling, thereby allowing direct pipette access from above^3^, and (ii) *Closed systems* in which the compartment is a channel that includes a ceiling^5–8^. In closed systems, unobstructed access to the worm is achieved by means of a dedicated pipette channel that joins the worm channel at a T-junction in the plane of the device.

Open and closed systems have reciprocal strengths and weaknesses. In open systems the injection pipette is positioned using a conventional micromanipulator. A key strength of open systems is that movement of the pipette is essentially unconstrained. It is also easy to change the injection pipette if clogged or broken (a frequent occurrence in DNA injections). In open systems, however, it is more difficult to fix the worm in position. These systems therefore rely on hydrogels or suction channels, both of which add complexity from the user’s point of view. Hydrogels require global temperature control across the plane of injection, whereas suction requires additional microfluidic channels and off-chip instrumentation to regulate and switch the vacuum.

Closed devices solve the problem of wet fixation by confining the worm in a close-fitting, fluid-filled injection channel. A key strength of closed systems is that they eliminate the complications of using hydrogels or vacuum. Another strength is that the injection channel facilitates worm handling. When the injection channel is connected to a worm reservoir at one end and a recovery chamber at the other end, the process of moving an injected worm out of the injection channel automatically brings the next worm into position for injection. On the other hand, extant closed devices have the weakness that the injection pipette must be carefully inserted through a long, narrow channel in the plane of the device, the height of which is on the order of the diameter of the worm, approximately 60 um^5–8^. This arrangement makes it inconvenient to exchange injection pipettes during a series of injections. In some systems, the injection pipette is integrated into the device during fabrication, such that when the pipette becomes clogged, the entire device must be replaced^7^. Furthermore, as the micropipette channel forms a junction with the worm channel, it introduces a pressure and fluid leak to the outside. To reduce this problem, some devices reduce leakage by adding a pressure activated control layer to compress the ceiling of the pipette channel^5,8^. Adding a control layer complicates the fabrication process and requires addition microfluidic channels plus off-chip instrumentation to regulate and switch the pressure. An alternative approach is to maintain the worm channel at ambient pressure. In this approach, instead of moving worms to the injection site by fluid flow, they are moved by harnessing their tendency to swim in the direction of an electric field^6^; however, suction is required to prevent the worm from swimming during the injection.

Despite the considerable inconvenience of conventional transgenesis methods in *C. elegans*, there appear to be no published reports citing the use of any of the six alternative methods. We suspect there are three main reasons for this apparent failure of adoption. First, relatively few *C. elegans* laboratories have the ability to fabricate complex, multi-layered devices in-house, and commercial foundry services for such devices would be prohibitively expensive. Second, the need to setup and maintain peripheral apparatus to control air pressure^3,5,8^ is a barrier to adoption for laboratories lacking a facility for instrumentation. Third, it is not clear that any of the alterative systems would be significantly easier to learn and utilize than the conventional method given, for example, the difficulty of fixing worms in open systems and of changing injection pipettes in closed systems.

In response to the challenges of injecting worms in a fluidic environment, we have developed a closed PDMS device that eliminates the need for a pipette channel. Instead, the worm is immobilized in an injection channel in which one sidewall is a thin PDMS membrane which the injection pipette penetrates to reach the worm. This device, called the *Poker Chip*, is monolithic, making it comparatively easy to fabricate. The device needs no peripheral apparatus except a hand-held syringe to move worms in and out of the injection channel. We anticipate that these advantages could lower barriers to adoption and accelerate basic, translational, and industrial research in this widely used model organism.

## Results and discussion

### Design strategy

A key design constraint was to lower barriers to adoption by making the device compatible with conventional injection setups. In such setups, the nematodes are mounted on a coverslip resting on the stage of an inverted compound microscope. The injection target is illuminated from above and viewed from below. The injection pipette, conventionally oriented at a low angle with respect to the plane of the stage, approaches the injection target from the side. Our design replicates this arrangement and requires no specialized equipment.

To facilitate fabrication, the Poker Chip has a minimal geometry and is monolithic (Fig. 1A). It comprises: (i) an injection channel optimized to restrain day-1 adults having a single row of eggs (*h* = 18 μm, *w* = 61 μm), (ii) an inlet port (1.5 mm diam.) that serves as a pre-injection reservoir large enough to contain tens of worms, and (iii) an outlet port (6 mm diam.) that serves as a post-injection reservoir. The height of the injection channel, being less than the diameter of an adult worm (50 um^9^), forces the animal to lie on its left or right side, the correct orientation for injection. The diameter of the outlet port is large enough to enable the user to withdraw injected worms with a Pasteur pipette.

**Fig 1.**
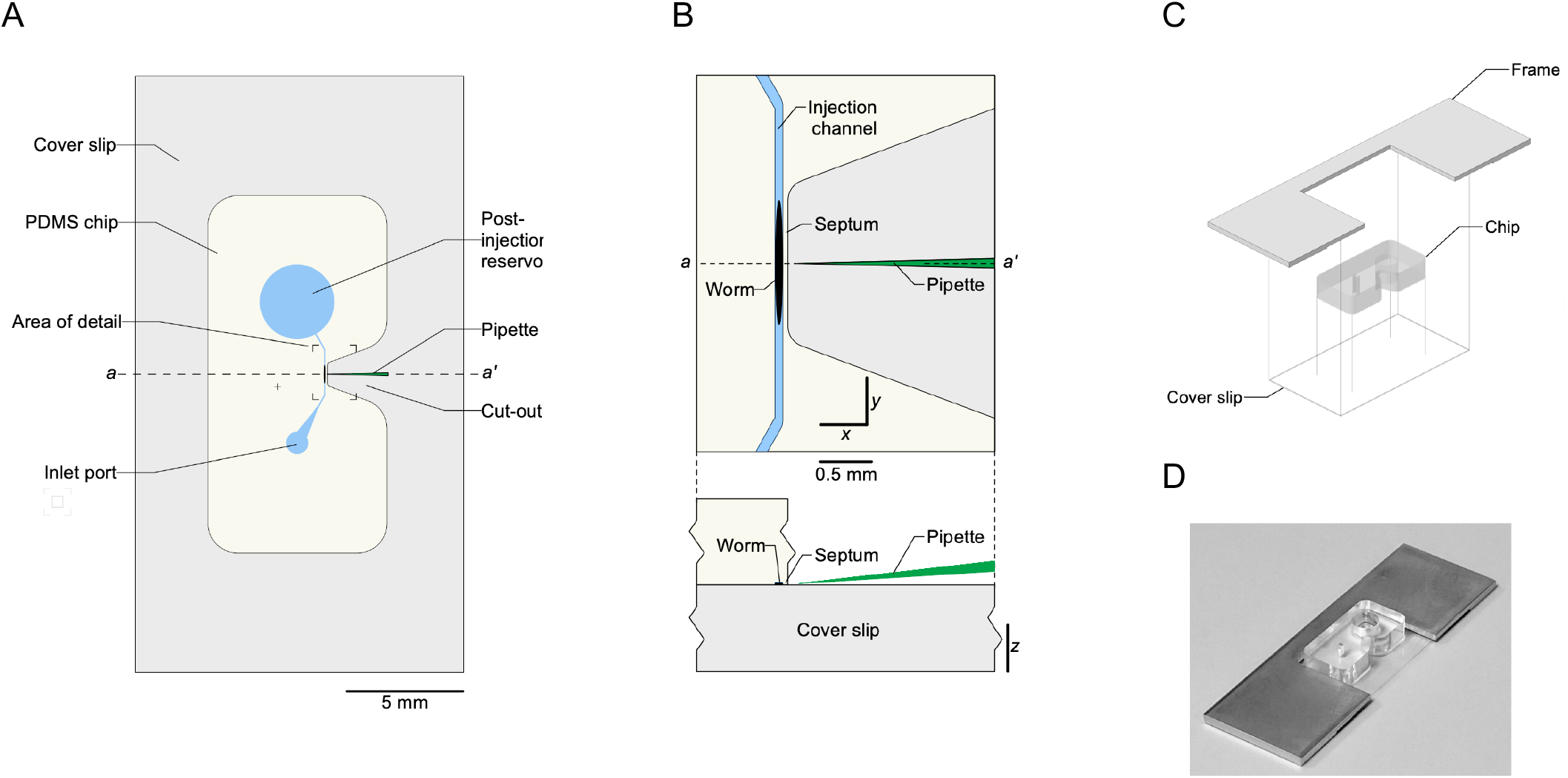
Layout and assembly of the injection chip. A. Top view. Colors: *light yellow*, PMDS; *blue*, fluid-filled features; *gray*, glass cover slip. B. Area of detail indicated in A. *Upper panel*, top view; *lower panel*, side view of *aa’* transect. C. Assembly of chip, cover slip, and frame. D. Photograph of the assembled device.

Opposite the center of the injection channel there is a nose-shaped cut-out that terminates at a point 40 um from the channel, forming a narrow septum through which the injection pipette is inserted to reach the worm (Fig. 1A,B). The walls of the cut-out are vertical so that the injection pipette can easily be lowered from above and there is no optical interference with visualization of the injection pipette. The top surface of the chip is optically flat to insure a clear image of the worm and the pipette tip. To close the channel, the chip is plasma bonded to a glass cover slip. The assembled device can be used in this form or it can be glued into an aluminum or acrylic frame to protect the cover slip from damage during handling (Fig. 1C).

The thickness of the septum is a critical dimension. It must be sufficiently thick to survive mold release without damage and to provide a mechanically strong bond with the cover. On the other hand, the septum must be thin enough to enable the user to align the tip of injection pipette with the vertical center of the worm’s gonad. The angle of the injection pipette with respect to the microscope stage (Fig. 1B) causes the pipette tip to follow a downwardly inclined trajectory as it passes through the septum. As a result, in order to hit the vertical center of the gonad, the user must choose an entry point on the septum just high enough to compensate for the vertical drop of the tip. A thinner septum facilitates this alignment process. We found that a thickness of 40 um was an optimal compromise for achieving mechanical strength and ease of alignment.

### Fabrication

Fabrication of the Poker Chip required development of a mold in which a macroscopic feature, the negative of the nose-shaped cut-out, is located with micron-order precision next to a microscopic feature, the negative of the injection channel. To overcome this challenge we designed an adjustable two-part brass mold (Fig. 2). The bottom plate of the mold includes a micromachined feature which forms the injection channel. The top plate includes a cavity that creates the overall outline of the chip, including the nose feature. To assemble the mold, the top plate is placed in contact with washers on the bottom plate. The plates are then screwed together by four screws (*1*), one in each corner. This step positions the nose feature approximately 100 um from the outside edge of the injection channel. The nose feature is then moved in close apposition to the channel by turning a screw (*2*) that presses against the back-side of the nose feature. The nose feature is then locked in place by a screw (*3*) that passes through a clearance hole in the the nose and threads into the bottom plate. The final position of the nose feature was found by an iterative processes of casting a PDMS positive, measuring the thickness of the septum in a calibrated photograph, and repositioning the nose feature as needed. The brass mold is limited to casting one PDMS positive at a time. To overcome this limitation, once the mold was correctly adjusted, we made multiplexed polyurethane molds^10^ by casting against a set of six PDMS positives.

**Fig 2.**
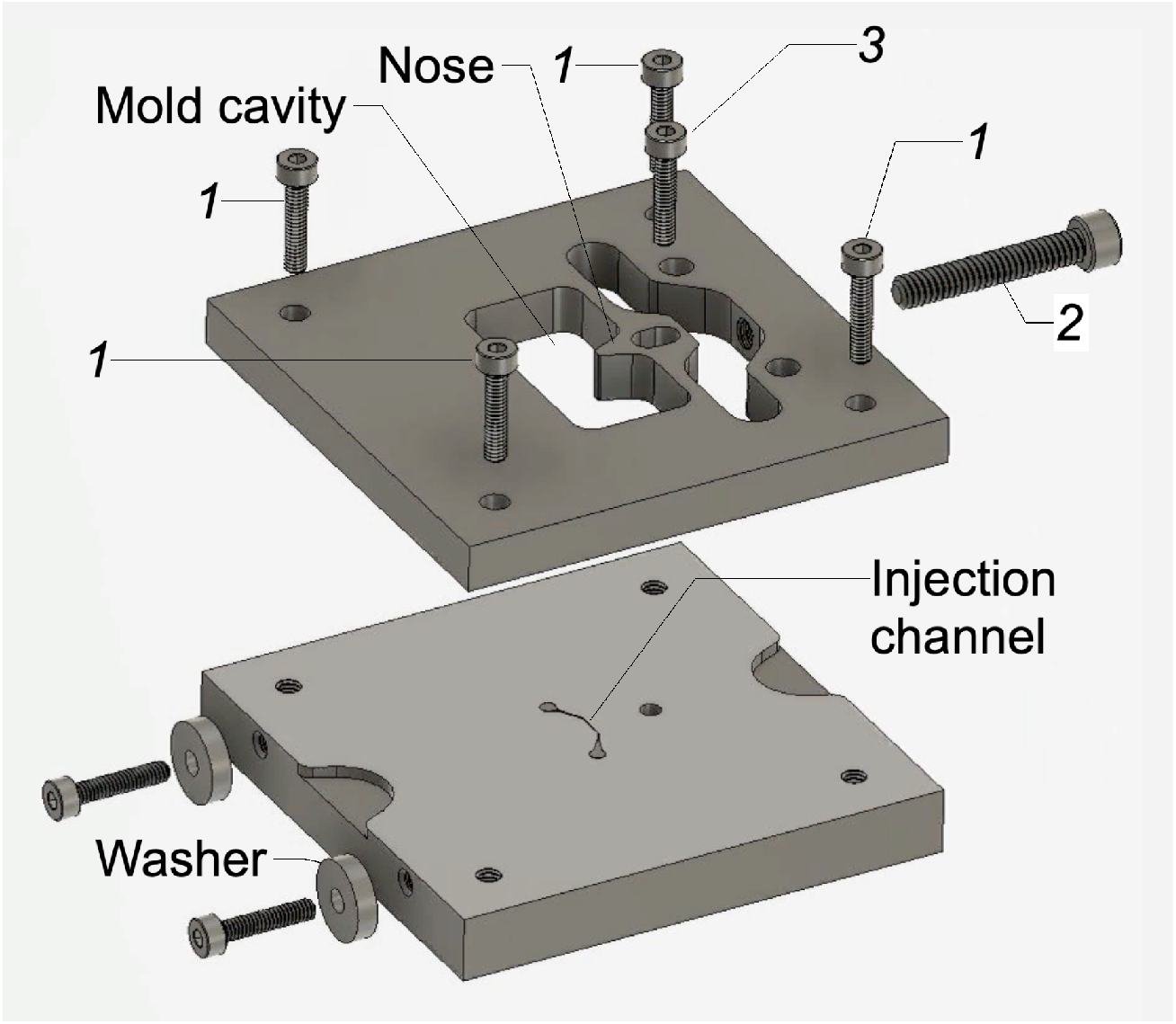
Injection chip mold. Top and bottom plates are joined by corner screws (*1*). The nose feature is positioned by turning a screw (*2*). The nose is locked into position by screw (*3*).

Chips were made by filling the cavities of the polyurethane mold with degassed PDMS pre-polymer (10:1 by weight). To ensure an optically flat top surface each cavity was slightly over-filled with PDMS, yielding a positive meniscus. A silanized glass slide, large enough to cover all six cavities, was then pressed down onto the mold, extruding excess PDMS. The glass slide was secured with a spring clamp and the assembled mold was cured for 3 hr at 65 C. To remove chips from the mold, we flooded the top of the mold with methanol and used a Teflon coated spatula to lift each chip from its cavity. Each chip was plasma bonded to a 24 mm x 60 mm No. 0 cover slip. The chip and cover slip assemblies where glued to the frame using spray adhesive (Elmer’s Spray Adhesive, Newell Brands, Westerville, OH, USA).

### Injection pipettes

The Poker Chip is designed be used with the same glass micropipettes used in conventional injections. To lower barriers to adoption, we used pipettes obtained from a commercial supplier of nematode DNA injection apparatus (TriTech, Los Angeles, CA, USA). The pipettes were formed from borosilicate capillary tubes (1.0 mm OD, 0.60 mm ID, with filament) using a PC-100 pipette puller (Narashige International USA, Amityville, NY, USA) (Fig 3A). The optimal pipette shape to minimize disruption of the septum is a long, gentle taper of about 2.8 deg starting approximately 140 um from the tip (Fig. 3B). When the pipette is in position within the gonad, the pipette diameter at the point of septum entry and exit is approximately 5.4 um and 1.8 um, respectively.

**Fig 3.**
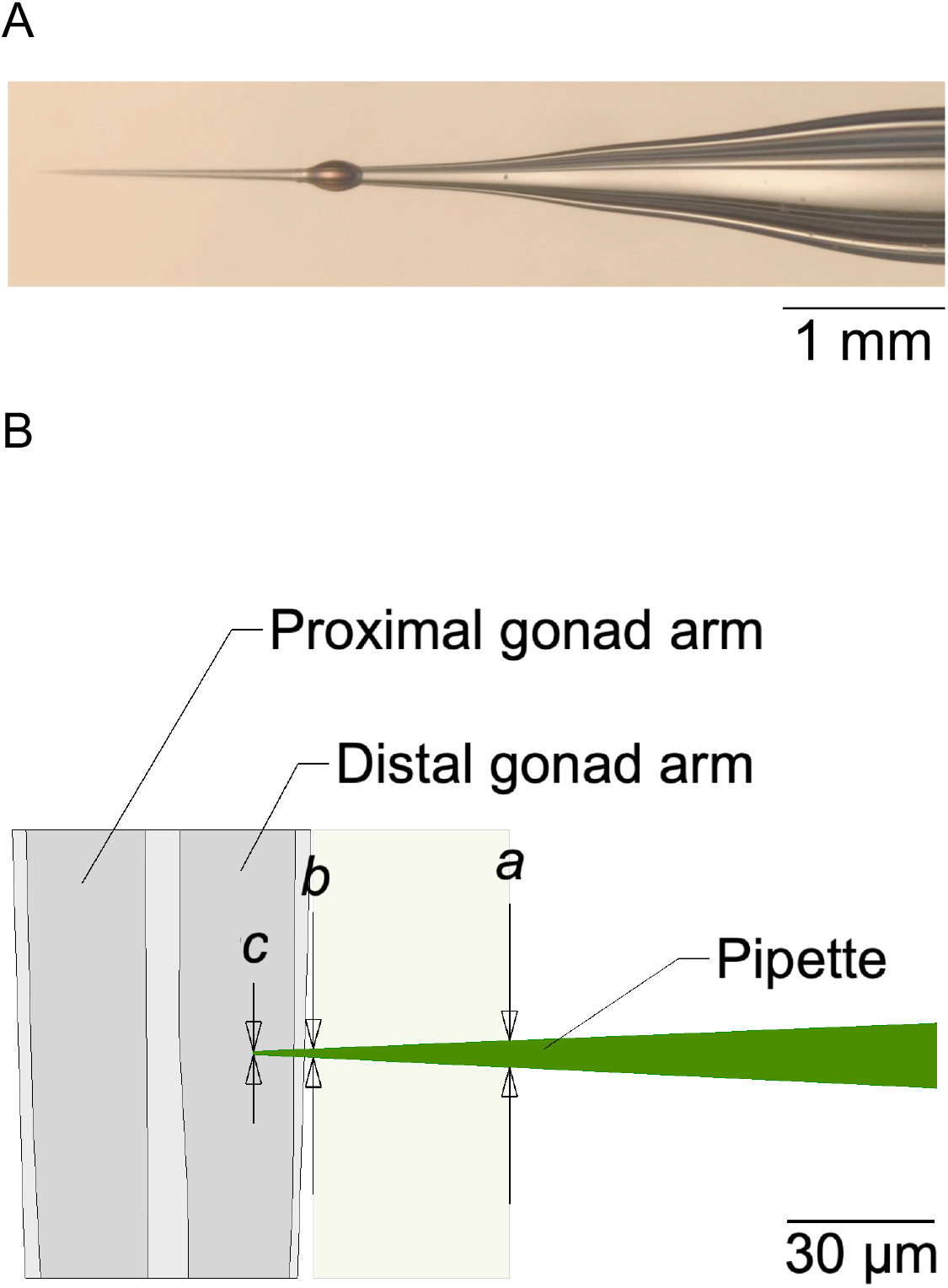
Injection pipette. A. Photomicrograph of an injection pipette after use. The ovoid object is a droplet of oil the remained on the pipette after use (see text, Method of use). B. Model of the tip shown in A. The diameter of the pipette at points *a, b*, and *c* is 5.4 μm, 1.8 μm, and ∼0.7 μm, respectively. Proximal and distal are defined relative to the position of worm’s vulva, which is on the ventral midline.

To test the ability of the injection pipette to penetrate the septum without clogging, we filled the injection channel with mineral oil and also placed a drop of oil at the base of the nose feature. The pipette was filled with a standard DNA injection mixture. When the pipette lumen was pressurized, ejection of fluid could be detected by the formation of a fluid droplet at the tip (Supplemental video 1). We found that droplets could be formed inside the injection channel after the tip passed through the septum. When the pipette was then withdrawn from the septum, droplet could still be formed within the nose feature, indicating that the pipette remained open after traversing the septum. Pressurizing the injection channel after withdrawing the pipette did not generate a fluid bubble in the nose feature, indicating that the septum resealed after penetration (not shown). Visual inspection of the pipette (40×) indicated that the pipette tip remained intact.

### Method of use

Preparation of worms. To obtain the tens of worms needed for a series of injections, a culture of synchronized worms is used. Worms are washed off the culture plate in 2 mL of standard M9 worm buffer (see Experimental), transferred to a 2 mL Eppendorf tube, and rinsed several times by pelleting and aspiration to remove debris and bacteria. After the final rinse, worms are concentrated into a loose pellet by allowing them to settle in the Eppendorf tube at room temperature or in a 4 C refrigerator.

Set-up. The chip is prepared by filling the injection channel with modified worm buffer (see Experimental), leaving room in the post-injection reservoir to accommodate injected worms and associated buffer. Worms are loaded into the inlet port by drawing 100 – 300 uL of fluid from the pellet using a micropipetter and ejecting the fluid into the port. An air-filled 10 mL syringe is fitted with 30-40 cm of M9filled flexible tubing; the free end of the tubing is inserted into the injection port. The chip is placed on the stage of a microscope fitted with Hoffman optics, with the middle of the septum centered in field of view. The chip is fixed in place using tape or plasticine modeling clay. A drop of hydrocarbon oil is placed at the end of the nose feature for testing pipette function (see below). Using the syringe, worms are moved from inlet port to the central region of the injection channel and then to the post-injection reservoir.

DNA injections. An injection pipette filled with injection mix is inserted into a conventional micropipette holder attached to a micromanipulator. The pipette is moved into position above the nose feature, then lowered until its tip can be seen in the oil. At this point injection pressure can be adjusted by raising it until the ejected fluid droplet is the desired size (see Supplemental video 1). A worm is moved into the injection channel using pressure from the syringe. The approximate vertical center of the gonad arm is found by focusing up and down through the arm. At this focal plane, the pipette tip is brought into focus just outside the septum by adjusting the vertical axis of the micromanipulator. The pipette is then raised slightly to compensate for its downward trajectory through the septum. At this point, the pipette is inserted through the septum and into the gonad, injection mix is injected, and the pipette is withdrawn from the worm (Supplemental video 2). In any given injection series, several practice injections into the first worm may be required to find the correct elevation of the pipette tip. However, once this elevation is found, further adjustments are generally not necessary. For convenience, the pipette can be withdrawn to a resting position in which the pipette tip is located at the horizontal center of septum, ready for the next worm to inject. Worms remain hydrated and apparently healthy for hours in the chip. It is therefore possible to switch pipettes and injection mixes many times without having to reload and remount the chip.

### Success rates

To compare the success rate of DNA injections using the Poker Chip to the success rate of the conventional method, we used the same injection mix in parallel sets of injections with 8 to 18 worms per set. The DNA marker in the mix was a semi-dominant allele of *rol-6*, which produces the so-called roller phenotype in which a cuticle defect causes worms to crawl in tight circles. This salient phenotype is widely used to positively identify transformed progeny of injected adults. In our tests, the same person performed all injections regardless of method. Worms were recovered by withdrawing them from the post-injection reservoir with a Pasteur pipette and placing them on a food-laden recovery plate. Twelve hours later, each worm was placed on its own recovery plate. After incubation for three days at 24 C, the progeny of each worm were scored for the roller phenotype. We computed survival rate as the fraction of injected worms that survived after being injected. We computed transformation rate as the fraction of injected worms whose progeny were rollers. We found that survival and transformation rates where statistically indistinguishable between the two methods (Fig. 4). We conclude that there are no obvious differences in these two key benchmarks of injection methods.

**Fig 4.**
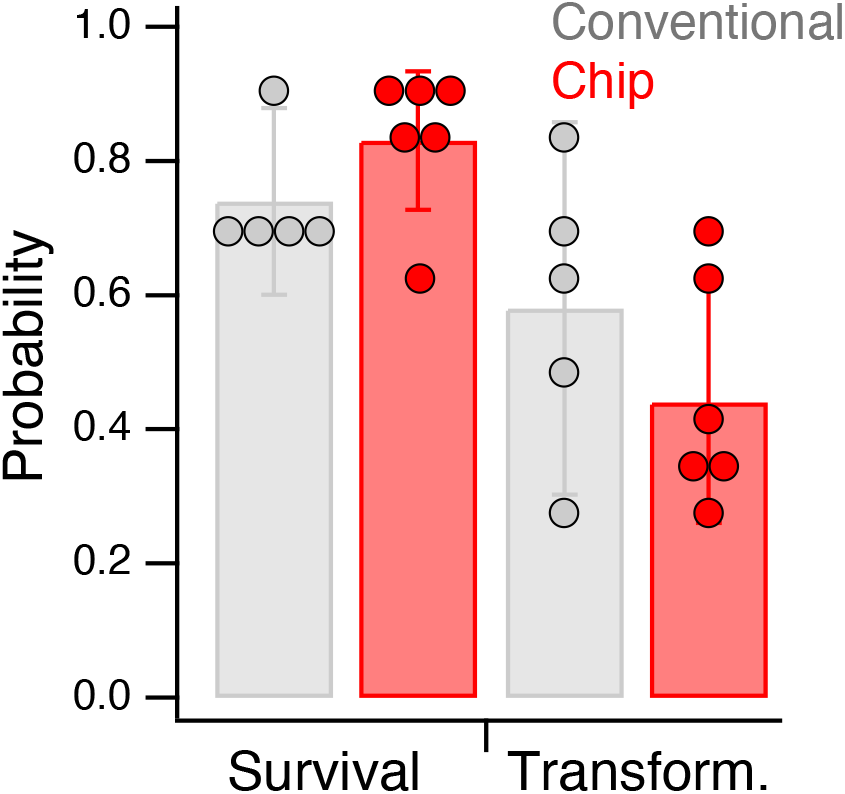
Comparison of survival and transformation rates in the conventional method and the injection chip method. Survival rate as a probability was computed as *N*_s_/*N*_i_, where *N*_s_and *N*_i_are number of worms alive 12 hours post-injection and number of worms injected, respectively. Transformation rate was computed as *N*_t_/*N*_i_, where *N*_t_is the number of worms that produced progeny having the roller phenotype. Statistics: survival rate, d.f. combined = 8.82, *t* = 1.37, *p* = 0.21; transformation rate, d.f. combined = 7.22, *t* = 1.20, *p* = 0.27. Error bars are 95% confidence intervals.

The standard injection site is the distal arm of the gonad, which is located dorsally. Thus, in the Poker Chip, only dorsal facing worms are injected; worms whose ventral side is facing the pipette are pushed directly into the post-injection reservoir (Supplemental video 2). The dorsoventral orientation of worms in the injection channel is random. On average, therefore, half population of worms in the post-injection reservoir receive an injection and half do not. Relative to the conventional method, the random orientation of worms essentially doubles the work of preparing culture plates and transferring individual worms to them for scoring progeny. One way to eliminate this problem is to mark injected worms by including a fluorescent dye in the injection mix such that only the fluorescent worms would be transferred to individual plates. To test the feasibility of this approach, we performed a series of conventional injections with either fluorescein or rhodamine added to the injection mix. For this series (see Methods), survival and transformation rates were at or near 100% regardless of dye concentration (Table 1). We conclude that high success rates can be obtained despite the presence of fluorescent dyes in the injection mix. This finding indicates that the efficiency of Poker Chip method could be improved by marking injected worms.

**Table 1.** Effect of fluorescent dyes on survival and transformation rate. Each row is an independent session of injections by the conventional microinjection method. The Concentration column shows the final dye concentration in the injected fluid. *N* = number of worms as indicated by the subscript. Parenthetical values indicate survival and transformation rate expressed as probability.

| Session | Dye | Concentration | $N_{\text{injected}}$ | $N_{\text{survived}} (p)$ | $N_{\text{transform.}} (p)$ |
| --- | --- | --- | --- | --- | --- |
| 1 | Fluorescein | 2.500 mg/mL | 12 | 12 (1.0) | 12 (1.0) |
| 2 | Fluorescein | 1.250 mg/mL | 12 | 12 (1.0) | 12 (1.0) |
| 3 | Rhodamine | 1.250 mg/mL | 10 | 10 (1.0) | 10 (1.0) |
| 4 | Rhodamine | 0.625 mg/mL | 10 | 9 (0.9) | 8 (0.8) |

### User experience

The conventional injection method is widely viewed as being difficult to master, requiring several weeks of training and practice. To test whether the Poker Chip method is easier to learn than the conventional method, we trained four volunteers to use the device. Volunteers ranged from zero to four days of experience with the conventional method. Volunteers were shown how to load the device with worms, mount it on the microscope stage, align the injection pipette with the nose feature of the device, move worms into the injection channel, penetrate the septum, inject fluid, and retract the pipette to the resting position. All four volunteers expressed confidence in being able to make injections after one day of training and practice.

## Conclusions

### Relation to previous devices

The Poker Chip combines the main strengths of open and closed systems for microinjection in *C. elegans*. As in open systems, the injection pipette is relatively unconstrained. It can be lowered from above and easily raised when the pipette must be changed. As in closed systems, the worm is immobilized by a close-fitting microchannel, eliminating the need for applying suction or hydrogels. Additionally, worms can be loaded in bulk, positioned semi-automatically, and then recovered in bulk after injection. The key innovation of the Poker Chip is elimination of a dedicated pipette channel. Such channels complicate the process of inserting and changing the pipette and they introduce a fluid leak which must be eliminated, usually by incorporating peripheral apparatus. In the present design, by contrast, the injection pipette is inserted through a thin PDMS septum which closes tightly around the pipette, eliminating leakage without peripheral apparatus.

### Barriers to adoption

A key goal of the research was to develop an injection chip with that would be adopted for routine use by many *C. elegans* laboratories. We focused on three main barriers to adoption. (1) *Fabrication*. Fabricating the Poker Chip is, of course, more involved than preparing the agarose pads used in the conventional method. However, compared to other injection chips the Poker Chip is comparatively simple to fabricate, as it comprises a single, monolithic PDMS block with only two ports. Indeed, fabricating the Poker Chip is no more difficult than fabricating the so-called olfactory chip used for neuronal calcium imaging in *C. elegans*^11^. The olfactory chip, which is also a single monolithic block, is likely one of the most widely adopted chip in *C. elegans* research^12^. (2) *Technology transfer*. The main barrier to transfer of a chip design from the originating laboratory to potential users in the *C. elegans* research community is the requirement for silicon SU-8 masters. Making silicon masters requires specialized equipment and masters have a limited lifetime. This problem can be ameliorated by transfer between labs of polyurethane molds such as those used here, instead of SU-8 masters. Finally, it is worth noting that the Poker Chip is reusable if cleaned and dried between injection sessions, reducing the number of new chips needed. (3) *Peripherals*. Like the conventional method, Poker Chip requires no peripheral apparatus (apart from a syringe and short length of plastic tubing). (4) *Ease of use*. In the conventional method, worms are positioned for injection by mounting them on the cover slip individually with a worm pick. With respect to the Poker Chip, the process of positioning worms for injection is simply a matter of applying gentle pressure with a hand-held syringe. Positioning the injection pipette relative to a worm in the Poker Chip is more challenging than in the conventional method because of the need to compensate for the downward trajectory of the tip. However, compensation is required only for the first several worms at the beginning of the injection session whereas, in the conventional method, the pipette must be repositioned for each worm, often requiring switching between low and high-powered objectives. On the other hand, the need to compensate for the downward trajectory of the tip in the Poker Chip could be eliminated simply by modifying the microscope stage to provide clearance for the injection pipette holder (5 – 10 mm diam.) so that the pipette can be oriented horizontally.

In deciding whether to adopt the Poker Chip, several additional considerations should be kept in mind. First, Poker Chip places tighter requirements on the shape of the injection pipette than the convention method does, in two respects. First, the injection chip performs best with pipettes that have a long gentle taper near the tip (Fig. 3). Pipettes with a short, steep taper can damage the septum, causing fluid leaks or shortening the lifetime of the chip. Pipette pullers that produce the optimal pipette shape are widely available (e.g., Narashige PC-100, Sutter P-97). Second, some practitioners of the conventional injection method prefer to break the pipette tip to facilitate penetration and increase the flow of fluid. Broken-tip pipettes are incompatible with the Poker Chip because they become clogged when pushed through the septum, presumably because they cut the PDMS, forming a plug within the pipette.

Another consideration is that the Poker Chip and its predecessors, in which the animals are restrained by a close-fitting microfluidic channel^5–8^, place tighter requirements on worm size. Worms that are too narrow are poorly restrained whereas worms that are too wide are difficult to move through the chip. We addressed this problem by synchronization (see Methods), but even synchronized worms vary in size. Moreover, the mean size of a synchronized population can be influenced by factors that are difficult to control, such as the number of eggs and amount of food deposited on culture plates. There are several ways this problem could be minimized: (i) create a range of chips with injection channels of having dimensions suited to different sizes of worms; (ii) adjust channel width by means of a void parallel to it which, when pressurized, pushes the channel wall opposite the septum inward; and (iii) load the chip with hand-picked worms of the appropriate size, as in the conventional method.

Glass micropipettes are the main tool for injecting DNA and other compounds in a wide variety of organisms, from cells^13–16^ and embryos^17–21^ to small animals such as nematodes^3–8^, Drosophila larvae^22^[ref2:16] and zebrafish larvae^23,24^. All injection applications require a means of stabilizing the target. Whereas numerous PDMS microdevices have been developed to fulfill this need^3,5–8,13–16,18,20,22,23^, all of them assume the need for an unobstructed pathway by which the injection pipette reaches the target. Creating and controlling such a pathway, and integrating the injection pipette within the device, appears to have been prevented broad adoption by biology researchers. Our demonstration that glass injection pipettes remain functional after being inserted through a PDMS membrane is significant because it effectively eliminates both constraints. The principle of direct penetration of elastomeric membranes is readily adaptable to microinjections in a wide range of organisms including cells, embryos, and other small animal models. It could therefore lead to a new generation of microinjection systems for basic, translational, industrial, and clinical applications.

## Experimental

### Device fabrication

Devices were cast in PDMS (Dow Corning Sylgard 184, Corning, NY, USA). Holes for ports and reservoirs were formed using biopsy punches of the appropriate diameter (in port, 1.5 mm; post-injection reservoir, 6 mm). PDMS castings were bonded to glass cover slips (Gold Seal Cover Glass, Fisher Scientific Co., Boston, MA, USA) after 60 s exposure to an oxidizing air plasma.

### Nematodes

The wild type (N2) strain of *C. elegans* was obtained from the *Caenorhabditis* Genetics Center at the University of Minnesota (St. Paul). Worms were grown at 20 C on NGM agar that had been previously seeded with the OP50 strain of *E. coli*. Worms were transferred to fresh plates of NGM with *E. coli* every 7–10 days to maintain healthy (not starved) stocks of worms. Synchronized populations of adult hermaphrodites were used throughout. Worms were synchronized by bleach synchronization according to establish procedures^25^ and allowed to grow for 72 hr at 20 C.

### Solutions

Standard M9 buffer was prepared by combining 3 g KH2 PO4, 6 g Na2 HPO4, 5 g NaCl and 1 mL of 1 M MgSO4, and adding H2O to 1 L. Modified M9 buffer was prepared by adjusting the osmolarity of standard M9 to 315 – 320 mOsm by addition of glycerol, to match the approximate osmolarity of the worm’s internal fluid.

The injection mix used in the experiment of Fig. 4 contained 40 ng/μL Super-rol plasmid DNA (InVivo Biosystems, Eugene, OR), 358 ng/uL Salmon Testes DNA (Sigma-Aldrich Inc., Saint Louis, MO), plus water to a final concentration of 100 ng/ul total DNA. The Super-rol plasmid is a double-stranded DNA that is similar to the widely used pRF4 plasmid containing the su100060 allele, but the Super-rol plasmid has been optimized for increased expression. Like pRF4, the Super-rol plasmid contains the R71C mutation in *rol-6* causing the rolling phenotype.

The injection mix stock used in the experiment of Table 1 contained CRISPR *dpy-10* co-injection marker, Cas9 (PNA-BIO, Newburry Park, CA), *dpy-10* gRNA (Synthego, Redwood City, CA), *dpy-10* oligo deoxynucleotide as donor-homologous single stranded DNA (Integrated DNA Technologies, Coralville, IA). Fluorescent dyes used were tetramethylrhodamine dextran neutral and fluorescein dextran anionic, (Thermo Fisher, Waltham, MA). Stock solutions of dyes contained 25 mg/mL 0.2 M KCl. Injection mix and dye solutions were combined to obtain the final dye concentrations indicted in Table 1.

### Pressure injection

Pipette pressure (80-100 psi) was regulated and switched using a digital microinjection pressure controller (MINJ-D, Tritech Research, Inc. Los Angeles, CA, USA).

## Acknowledgements

Supported by NIH grant number GM129576.

## Figures and tables

**Supplemental video 1.**
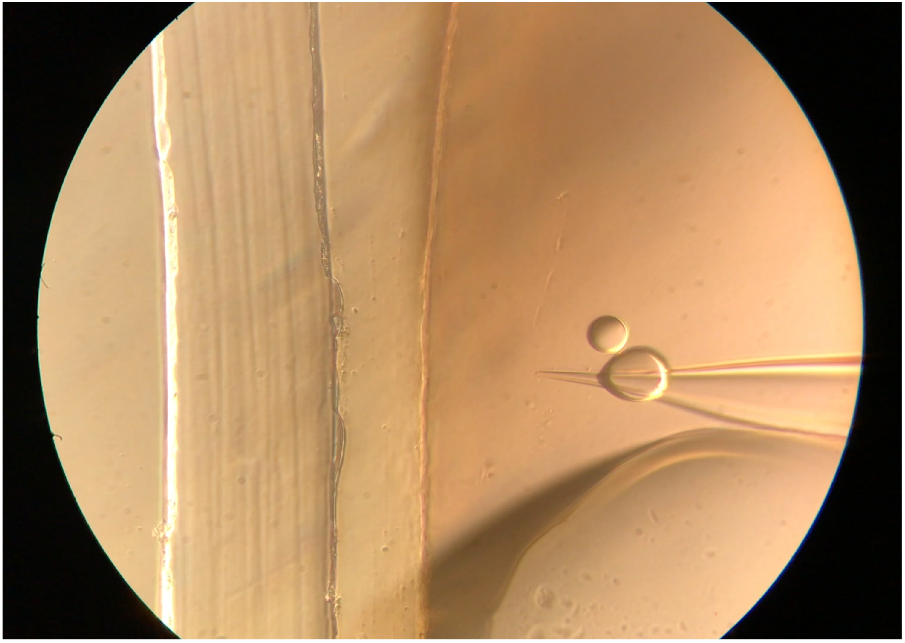
Demonstration that the pipette in not clogged by passage through the septum (4× speed). The injection channel is oriented vertically in the field of view and the injection pipette enters from the right. The channel is filled with mineral oil and a drop of oil is placed to the right of the septum. The injection pipette is filled with DNA injection mix. Pressure pulses cause water droplets to form in the oil. Download link: https://www.dropbox.com/s/ey83wjz8u7afg4m/Supplemental-video-4x-1.mp4?dl=0

**Supplemental video 2.**
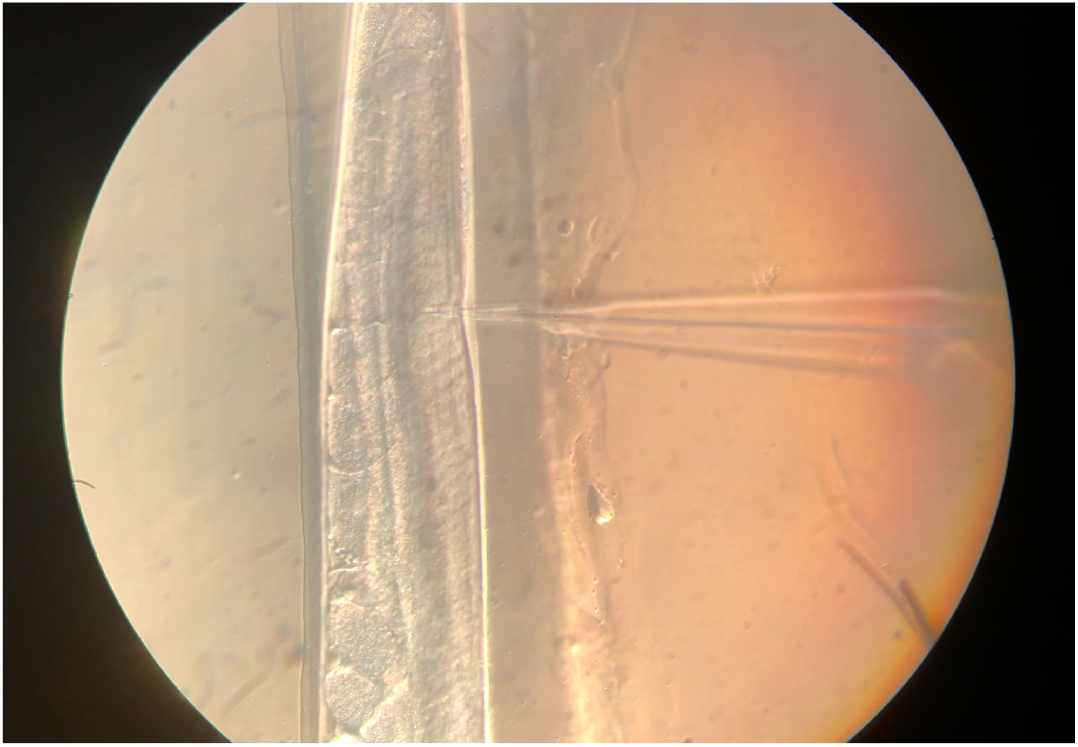
DNA injections using the poker chip (4× speed). The injection channel is oriented vertically in the field of view and the injection pipette enters from the right. Worms in which the distal gonad arms are on same side as the pipette are injected; worms in the opposite orientation are skipped. Download link: https://www.dropbox.com/s/rkrxjjaklt7h572/Supplemental-video-4x-2.mp4?dl=0

